# Developmental timing of *Drosophila pachea* pupae is robust to temperature changes

**DOI:** 10.1101/2021.08.25.457619

**Authors:** Bénédicte M. Lefèvre, Stecy Mienanzambi, Michael Lang

## Abstract

Rearing temperature is correlated with the timing and speed of development in a wide range of poikiloterm animals that do not regulate their body temperature. However, exceptions exist, especially in species that live in environments with high temperature extremes or oscillations. *Drosophila pachea* is endemic to the Sonoran desert in Mexico, in which temperatures and temperature variations are extreme. We wondered if the developmental timing in *D. pachea* may be sensitive to differing rearing temperatures or if it remains constant. We determined the overall timing of the *Drosophila pachea* life-cycle at different temperatures. The duration of pupal development was similar at 25°C, 29°C and 32°C, although the relative progress differed at particular stages. Thus, *D. pachea* may have evolved mechanisms to buffer temperature effects on developmental speed, potentially to ensure proper development and individual’s fitness in desert climate conditions.

## 1. Introduction

Poikilotherms animals do not regulate their body temperature contrary to homeotherms (Precht et al., 1973) and are sensitive to environmental temperature. Environmental temperature in turn affects their metabolism (Hazel and Prosser, 1974). In particular, it seems widespread that developmental speed increases with rearing temperature in poikilothermic species (Abril et al., 2010; Asano and Cassill, 2012; Hrs-Brenko et al., 1977; Ikemoto, 2005; Manoj Nair and Appukuttan, 2003; Nishizaki et al., 2015; Pechenik et al., 1990; Porter, 1988; Sharpe and DeMichele, 1977; Vélez and Epifanio, 1981), including various Drosophila species (David and Clavel, 1966; James and Partridge, 1995; Kuntz and Eisen, 2014; Powsner, 1935). This phenomenon is proposed to be due to thermodynamics of enzymes responsible for biochemical reactions underlying developmental processes (Crapse et al., 2021; Ikemoto, 2005; Schoolfield et al., 1981; Sharpe and DeMichele, 1977). Thermal-stress can accelerate development and has been shown to result in an increase of developmental instability (Kristensen et al., 2003; Nishizaki et al., 2015; Polak and Tomkins, 2013), measured as deviations of an individual’s character from the average phenotype in the population under the same conditions (Palmer, 1994; Zakharov, 1992). This may result in a decreased individual’s survival and reproductive fitness. In contrast, a slow development may potentially lead to an increased risk of predation at vulnerable stages, such as immobile pupae in holometabolous insects (Ballman et al., 2017; Borne et al., 2021; Hennessey, 1997; Thomas, 1993; Urbaneja et al., 2006). Furthermore, a variable timing of development among individuals of a same species might induce intraspecific competition (Amarasekare and Coutinho, 2014; Frogner, 1980) as individuals developing faster may reproduce sooner and for a longer period compared to those developing more slowly. Different mechanisms have been found to regulate developmental timing. The so-called heterochronic miRNAs, such as *let-7* and *miR-125*, were originally discovered in *Caenorhabditis elegans* (Rhabditida: Rhabditidae)(Ambros, 2011; Ambros and Horvitz, 1984). These miRNAs are conserved in a wide range of species, such as *Drosophila melanogaster* (Diptera : Drosophilidae)(Caygill and Johnston, 2008) or *Danio rerio* (Cypriniformes: Cyprinidae)(Ouchi et al., 2014), as well as in mammals and plants (Ambros, 2011). They act at post-transcriptional level to regulate cellular mRNA levels, and have been found to control the developmental timing, cell fate and cell differentiation. Hormones are also known to be important regulators of developmental timing. I n *D. melanogaster*, each of the developmental transitions are regulated by ecdysone pulses, and premature transition from larva to pupa with respect to food conditions or starvation is prevented by juvenile hormone (Riddiford, 1994; Riddiford and Ashburner, 1991). Thus, developmental timing might be regulated to reach an optimal duration with respect to outer environmental factors.

More than 1500 described species of the genus *Drosophila* (Bächli et al., 2021; O’Grady and DeSalle, 2018) occupy a wide range of habitats with various climatic conditions (Markow and O’Grady, 2008). A dozen of species have been reported to be cosmopolitan species (Markow and O’Grady, 2008, 2005), such as *Drosophila melanogaster* (David and Capy, 1988; Li and Stephan, 2006) that potentially dispersed with humans from Africa around the globe (Mansourian et al., 2018). These species may be generalists but were also found to be locally adapted to diverse environments (Kapun et al., 2020; Markow and O’Grady, 2008). In contrast, the vast majority of species are restricted to certain continental ranges or are endemic to a specific geographic region that encompasses a unique habitat with specific food and climate conditions (Markow and O’Grady, 2008, 2005). Because of their inability to disperse outside their habitat, these endemic species may have evolved temperature-buffering mechanisms to ensure a constant developmental timing under variable temperature conditions.

*Drosophila pachea* (Diptera : Drosophilidae) is endemic to the Sonoran desert in Mexico and is an obligate specialist on decayed parts, or rot-pockets, of its single host plant, the Senita cactus (*Lophocereus schottii*) (Gibbs et al., 2003; Heed and Kircher, 1965; Lang et al., 2012; Markow and O’Grady, 2005). The micro-climate of the rot-pockets encompasses important changes of temperature all along the year, with a recorded maximum variation from 5°C to 42°C within 24 h (Gibbs et al., 2003). Living in an environment with large daily and annual temperature changes may require a certain temperature robustness with respect to developmental processes in poikiloterm species. We wondered if the developmental timing in *D. pachea* may be sensitive to differing rearing temperatures. To test this, we first determined the overall timing of the *Drosophila pachea* life-cycle. Then, we focussed on pupal development at four different rearing temperatures to investigate differences in the pupal timing. Finally, we compared these durations across closely related sister species *Drosophila acanthoptera* (Diptera : Drosophilidae) and *Drosophila nannoptera* (Diptera : Drosophilidae) to investigate potential species-specific developmental timing differences.

## 2. Materials and methods

### 2.1. Drosophila stock maintenance

Drosophila stocks were retrieved from the San Diego Drosophila Species Stock Center (now The National Drosophila Species Stock Center, College of Agriculture and Life Science, Cornell University, USA). The *D. pachea* stock 15090-1698.01 was established in 1997 from individuals caught in Arizona, USA. The *D. nannoptera* stocks 15090-1692.00 and 15090-1693.12 were established in 1992 from individuals caught in Oaxaca, Mexico. The *D. acanthoptera* stock 15090-1693.00 was established in 1976 from individuals caught in Oaxaca, Mexico (UCSC Drosophila species stock center San Diego, now The National Drosophila Species stock center, Cornell University). These stocks have been kept in good conditions at 25°C in our laboratory since 2012.

Flies were maintained in transparent plastic vials (25 × 95 mm, Dutscher) containing about 10 mL of standard Drosophila medium. This medium was composed of 66.6 g/L of cornmeal, 60 g/L of brewer’s yeast, 8.6 g/L of agar, 5 g/L of methyl-4-hydroxybenzoate and 2.5% v/v ethanol (standard food). We added 40 μL of 5 mg/mL of 7-dehydrocholesterol (7DHC) (Sigma, reference 30800-5G-F) dissolved in ethanol into the food for *D. pachea*, as this species need this sterol for proper development (Heed and Kircher, 1965; Lang et al., 2012; Warren et al., 2001) (standard *D. pachea* food). As a pupariation support, a piece of paper sheet (1 cm x 4 cm, BenchGuard) was added to each vial. Stocks were kept at 25°C or 29°C at a 12 h light:12 h dark photoperiodic cycle with a 30 min transition between light (1080 lm) and dark (0 lm).

### 2.2. Cohort synchronisation of *D. pachea* embryos and time-lapse recording of embryonic development

For collection of cohorts of synchronised embryos, about 250-500 adult flies were transferred into a 9 × 6 cm plastic cylinder, closed by a net on the top and by a 5.5 cm diameter petri-dish lid at the bottom. The petri-dish contained grape juice agar (24.0 g/L agar, 26.4 g/L saccharose, 20% grape juice, 50% distilled water, 12% Tegosept [1.1 g/mL in ethanol] (Dutscher), 4% 7-DHC (Sigma)) and 50-200 μL fresh baker’s yeast as food source and egg laying substrate on top. These plates are named hereinafter “food plates”. Female flies were let to lay eggs on the food plates for 1 h - 2 h (1 h to examine embryos and 2 h to synchronise larvae). Then, eggs were retrieved from food plates by filtering the yeast paste through a 100 μm nylon mesh (BS, Falcon 352360).

For time-lapse imaging the chorion of embryos was removed by a 90 sec incubation of the embryo-containing filter in 1.3% bleach (BEC Javel) under constant agitation until about half of the embryos were floating at the surface of the bleach bath. Embryos were extensively rinsed with tap water for at least 30 sec. Dechorionated embryos were then gently glued on a cover slip (ThermoFisher) coated with Tesa glue. For coating, 50 cm TESA tape was transferred into 25 mL n-heptane (Merck) and glue was let to dissolve overnight at room temperature. A total of 15 µL of dissolved glues was finally pipetted onto a cover slip to form a 5 × 20 mm rectangular stripe and n-heptane was let to evaporate. Embryos were covered with 40 μL of Voltalef 10S halocarbon oil (VWR) to avoid desiccation. Live-imaging was immediately launched inside a temperature and humidity controlled chamber at 25°C ± 0.1°C and 80% ± 1% humidity (Lang and Orgogozo, 2012; Lefèvre et al., 2021; Rhebergen et al., 2016). Time-lapse acquisition was performed at an acquisition rate of 1 picture every 7.5 sec using a digital camera (Conrad 9-Megapixel USB digital microscope camera) and Cheese software, version 3.18.1, on a computer with an ubuntu 16.04 linux operating system. Movies were assembled with avconv (libav-tools).

In *Drosophila melanogaster* and closely related species, females were reported to hold fertilized eggs inside the reproductive tract for >12 hours (Markow et al., 2009), which could explain the variation observed in our experiments with *D. pachea*. Therefore, we monitored egg retention in this species by examining dechorionated eggs from a 1 h egg-laying period. We found that all observed embryos (n=52) were early embryos at the syncytial blastoderm stage (Wieschaus and Nüsslein-Volhard, 1986) and egg retention was not observed. Out of 28 embryos monitored, 12 (43%) pursued their development until hatching while the others did not develop at all (Movie S1, Dataset S1). Such mortality has been reported previously (Jefferson, 1977; Pitnick, 1993) but potentially also dependent on the above-mentioned bleach treatment. The embryos that died during the experiment were excluded from analysis. Furthermore, the duration of hatching, which is the last stage of embryonic development, has been shown to be more variable in comparison to the other embryonic stages in various Drosophila species (Chong et al., 2018; Kuntz and Eisen, 2014). We thus measured both the total embryonic duration, from collection up to larva hatching and the embryonic duration up to the trachea gas filling stage, which precedes the hatching stage (Dataset S1).

### 2.3. Cohort synchronisation of larvae, dissection and imaging of larval mouth hooks

In order to collect cohorts of larvae at a synchronous developmental stage, we first collected embryos from a 2 h egg laying interval (see above) that were placed on a food plate together with fresh yeast. Freshly hatched larvae were retrieved from the yeast paste with fine forceps (Dumont #5, Fine Science Tool) or by filtering the yeast through a nylon mesh (see above). Larvae were transferred into vials containing standard *Drosophila pachea* food and were examined once a day until all larvae had turned into pupae (Dataset S2).

For imaging of the larval teeth, entire larvae were mounted in 20 μL dimethyl-hydantoin formaldehyde (DMHF) medium (Entomopraxis) beneath a cover slip (0.17 mm ± 0.01 mm thick, ThermoScientific), which was gently pressed against the microscope slide (ThermoScientific) to orient larval teeth in a flat, lateral orientation to the microscope objective. Larval teeth were imaged at 100 or 400 fold magnification in bright field illumination (Strasburger, 1935) using the microscope IX83 (Olympus). The instar stage of each dissected individual was determined based on tooth morphology (Strasburger, 1935) (Figure S1).

### 2.4. Measurement of the duration of puparium formation in *D. pachea*

The precise duration of puparium formation was characterized by monitoring nine *D. pachea* pupariating larvae by time-lapse imaging. Larvae at the third instar stage and third instar wandering stage were collected from the *D. pachea* stock and were transferred into fresh *D. pachea* standard medium, inside a 5 cm diameter petri-dish and a piece of 1 cm x 4 cm paper sheet (BenchGuard). The dish was then placed into the temperature and humidity controlled chamber at 25°C ± 0.1°C and 80% ± 1% humidity, as previously described. Time-lapse acquisition was performed for about 72 h as previously described for embryonic timing characterization. The duration of the white puparium stage was measured from the moment when the larva had everted the anterior spiracles and had stopped moving until the moment when the pupal case had turned brown.

### 2.5. Characterization of developmental timing in pupae

The developmental duration of *D. pachea, D. nannoptera* and *D. acanthoptera* was examined by observation of pupae at different time points after puparium formation (APF). Synchronised pupae were obtained from each species by collecting so-called “white pupae” that had just formed the puparium (Dataset S3). Specimens were collected with a wet brush directly from stock vials. Individuals of the same cohort were placed onto moist Kimtech tissue (Kimberly-Clark) inside a 5 cm diameter petri dish. Petri dishes with pupae were kept at 22°C, 25°C, 29°C, or 32°C inside plastic boxes, which also contained wet tissue paper. Specimens analyzed at 22°C and 32°C were taken from a stock at 25°C at the stage of puparium formation and subsequently incubated at the desired temperature. A group of pupae resulting from a single collection event was considered as a synchronised cohort. Developmental progress of synchronised cohorts (Table 1, Figure S2) was examined at various time points by visual examination of the pupae using a stereomicroscope VisiScope SZB 200 (VWR) (Dataset S4). Developmental stages were assigned according to morphological markers defined for *D. melanogaster* b y Bainbridge and Bownes (1981) (Table 2). The markers used to characterize stages 8 to 12 (eye, wing or body pigmentation, Table 2) were not convenient for the characterization by direct observation of *D. acanthoptera* pupae as these flies develop black eyes, as opposed to most other Drosophila species that have red eyes. In addition, *D. acanthoptera* is generally less pigmented compared to *D. pachea* and *D. nannoptera* (Pitnick and Heed, 1994) and pigmentation changes were not easily detectable through the pupal case. Therefore, we additionally carried out time-lapse imaging of one cohort with five *D. acanthoptera* pupae to investigate the developmental durations of stages 8-12. The anterior part of the pupal case was removed, letting the head and the anterior part of the thorax visible. Image acquisition was done at 25°C ± 0.1°C and 80% ± 1% humidity, as previously described. Time-lapse acquisition was performed as previously described and recorded with the VLC media player, version 3.0 at an acquisition rate of 1 picture every 13:02 min. Two pupae died during acquisition and were excluded from the analysis (Movie S2).

**Table 1:**
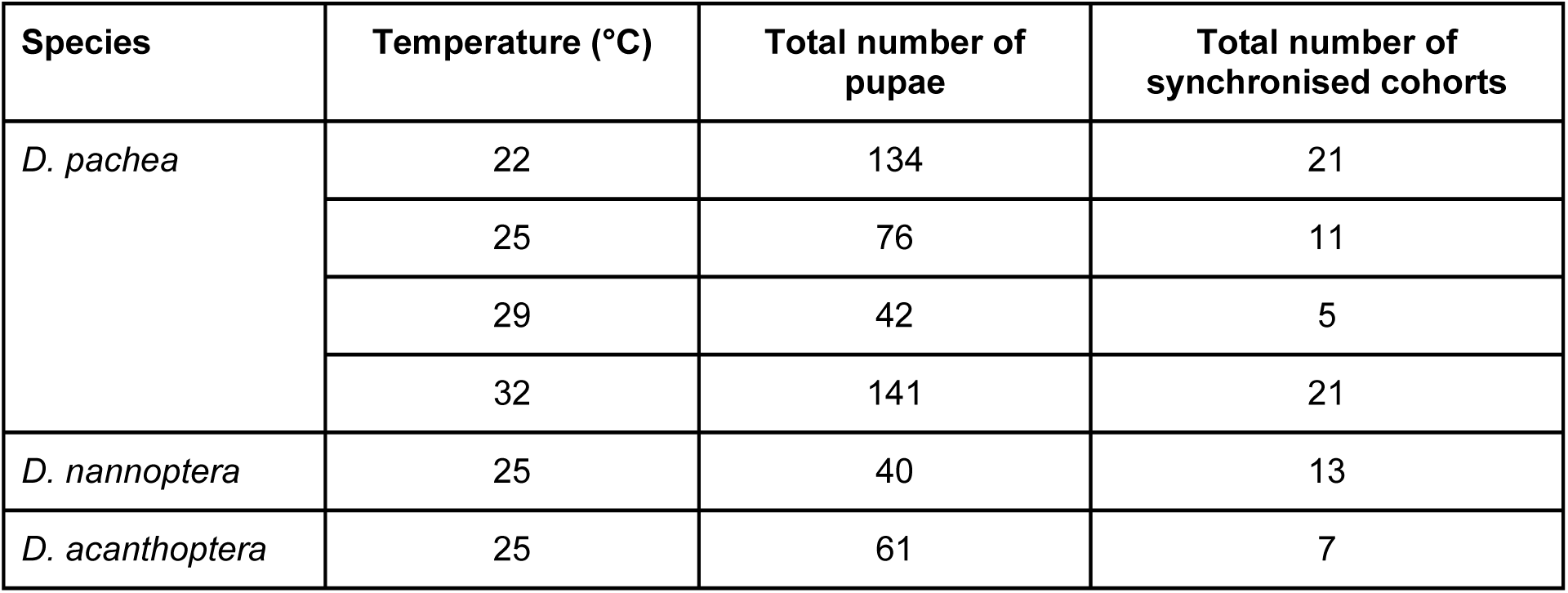
Total numbers of pupae and synchronised cohorts used in for pupal timing characterization.

**Table 2:** Summary of morphological markers used to stage pupae, according to Bainbridges and Bownes (1981)

| Pupal stage | Description |
| --- | --- |
| 1 | Pupariation: extremity of trachea are everted, pupa does not move anymore. |
| 2 | Clear, white pupa. |
| 3 | Light pigmentation, dorsal trachea still visible. |
| 4 | Bubbles appear, dorsal trachea is not visible anymore, light pigmentation of the body. |
| 5 | Cranial extremity is retracted, distal extremity of wings appeared. |
| 6 | Yellow body is visible. |
| 7 | Pharate adult, eyes are not yet pigmented. |
| 8 | Eye discs become a bit yellower compared to the rest of the body. |
| 9 | Orange eyes, transparent wings are visible. |
| 10 | Deep red eyes. |
| 11 | Bristles are visible on the thorax. |
| 12 | Wings are clear grey, clear pigmentation of the body. |
| 13 | Wings are completely black, grey pigmentation appears on the body. |
| 14 | Meconium appeared under the form of a dark spot visible through the abdomen. |
| 15 | Eclosion of the adult. |

### 2.6. Data analysis

Data was manually entered into spreadsheets (Datasets S1, S2, S3 and S4) and analysis was performed in R version 3.6 (R Core Team, 2014). Ages expressed in hours after pupa formation were automatically calculated with respect to the time point of white pupa collection.

## 3. Results

### 3.1. *D. pachea* embryonic and larval development at 25°C last for about 33 h and 216 h, respectively

W e roughly examined the duration of embryonic and larval development in *D. pachea* at 25°C. The average duration of the total embryonic development in *D. pachea* at 25°C, until hatching of the larva was 32 h 48 min ± 1 h 13 min (mean ± standard deviation ; n = 12) (Figure 1, Movie S1). Embryonic development up to the trachea gas filling stage (see Material and Methods for details) was estimated to be 26 h 48 min ± 1 h 13 min (mean ± standard deviation ; n = 12) (Movie S1). These durations appeared to be longer in *D. pachea* compared to those reported for various other Drosophila species, such as *Drosophila melanogaster, Drosophila simulans, Drosophila sechellia, Drosophila yakuba, Drosophila pseudoobscura, Drosophila mojavensis* (Figure 2) (David and Clavel, 1966; Kuntz and Eisen, 2014; Powsner, 1935).

**Figure 1:**
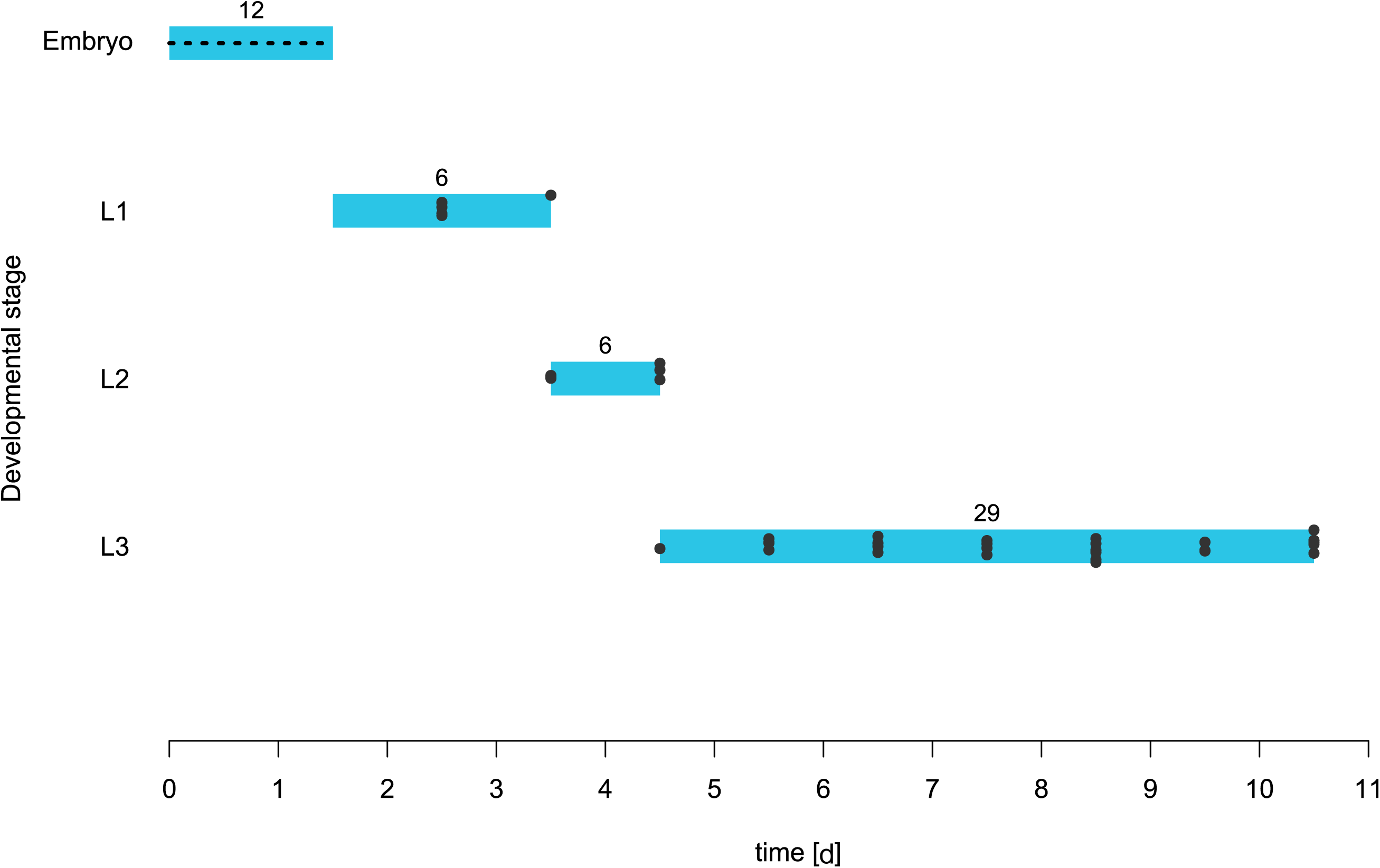
Timing of the embryonic and larval stages in *D. pachea* at 25°C. Embryo duration represents the time from egg laying to the hatching of the larva, based on time-lapse imaging (dotted line). Larval stages were determined based on mouth hook morphology of dissected larvae from synchronized cohorts, according to Strasburger (1935). Black dots indicate single observations (Dataset S2). Numbers correspond to the number of individuals observed at each stage.

**Figure 2:**
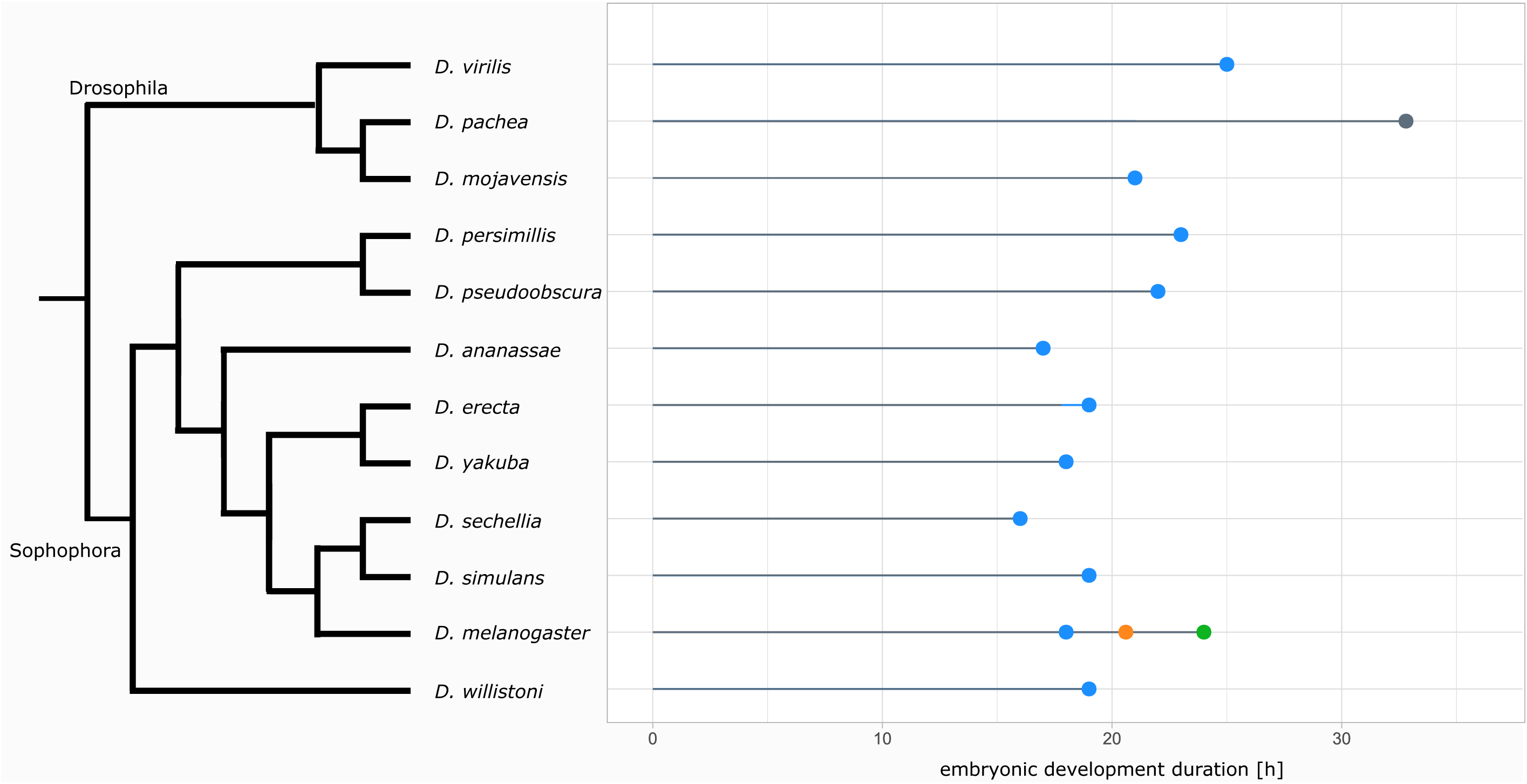
Durations of the embryonic development in various *Drosophila* species at 25°C. The duration of total embryonic development of *D. pachea* (grey) was established based on time-lapse imaging. The data for the species other than *D. pachea* were extracted from: blue: Kuntz and Eisen, 2014 (duration up to the trachea filling stage, at 25°C), yellow: David and Clavel, 1966 (total embryonic development, at 25°C) and green: Powsner, 1935 (total embryonic development, at 25°C). The data used to establish the cladogram was extracted from Yassin (2013) and Lang et al. (2014).

The total duration of *D. pachea* larval development on standard *D. pachea* food at 25°C was approximately 9 days (∼216 h). The duration of the first and second instar larva were about 2 days each and the third instar stage lasted for about 5 days (Figure 1). In *D. melanogaster*, the total duration of the larval stage was about 5 days for larvae reared on optimal food at 25°C, the first and second instars lasting for 1 day each, and the third instar for three days, according to Strasburger, (1935). The larval development of *D. pachea* appeared thus to be longer compared to those of *D. melanogaster* at 25°C.

### 3.2. Similar durations of pupal development in *D. pachea* at 25°C, 29°C and 32°C

The duration of larval development appears to be sensitive to various environmental factors, such as diet (Matzkin et al., 2011), crowding, or access to food (Vijendravarma et al., 2013). Since pupal development is apparently less affected by such factors, we focussed on the pupal stage to investigate the effect of the rearing temperature on timing of development in *D. pachea*. We evaluated pupal developmental progress at four temperatures: 22°C, 25, 29°C, and 32°C. Preliminary tests revealed that rearing of *D. pachea* at temperatures lower than 25°C is prolonged which favors the accumulation of bacterial infections in the food and decreased survival of the flies. At 34°C, *D. pachea* individuals died within a few days and at 32°C flies survived but did not reproduce. Since we could not cultivate *D. pachea* at the extreme temperatures of 22°C and 32°C, individuals were selected at the stage of puparium formation in a stock at 25°C and incubated at either temperature.

At 25°C - 32°C, *D. pachea* pupae reached the pharate adult stage in less than 55 h but timing was prolonged at 22°C (Figure 3A-E). However, pupal development was accelerated at 29°C and 32°C between stages 8 and 13 (beginning of eye pigmentation until the end of body and wing pigmentation) compared to development at 25°C (Figure 3A-E). In addition, development was consistently slower at 22°C compared to 25°C. However, stages 14 and 15 required more time at 29°C and 32°C with respect to developmental progress at 25°C and resulted in a similar overall duration of about 100 - 145 h. Only at 22°C, development was globally slower and adults emerged later, between 150 - 190 h. Thus, in *D. pachea* the rearing temperature influences the relative progress of pupal development at particular stages. While pupal development is slowed-down at temperatures below 25°C, the overall duration appears to be similar at higher temperatures.

**Figure 3:**
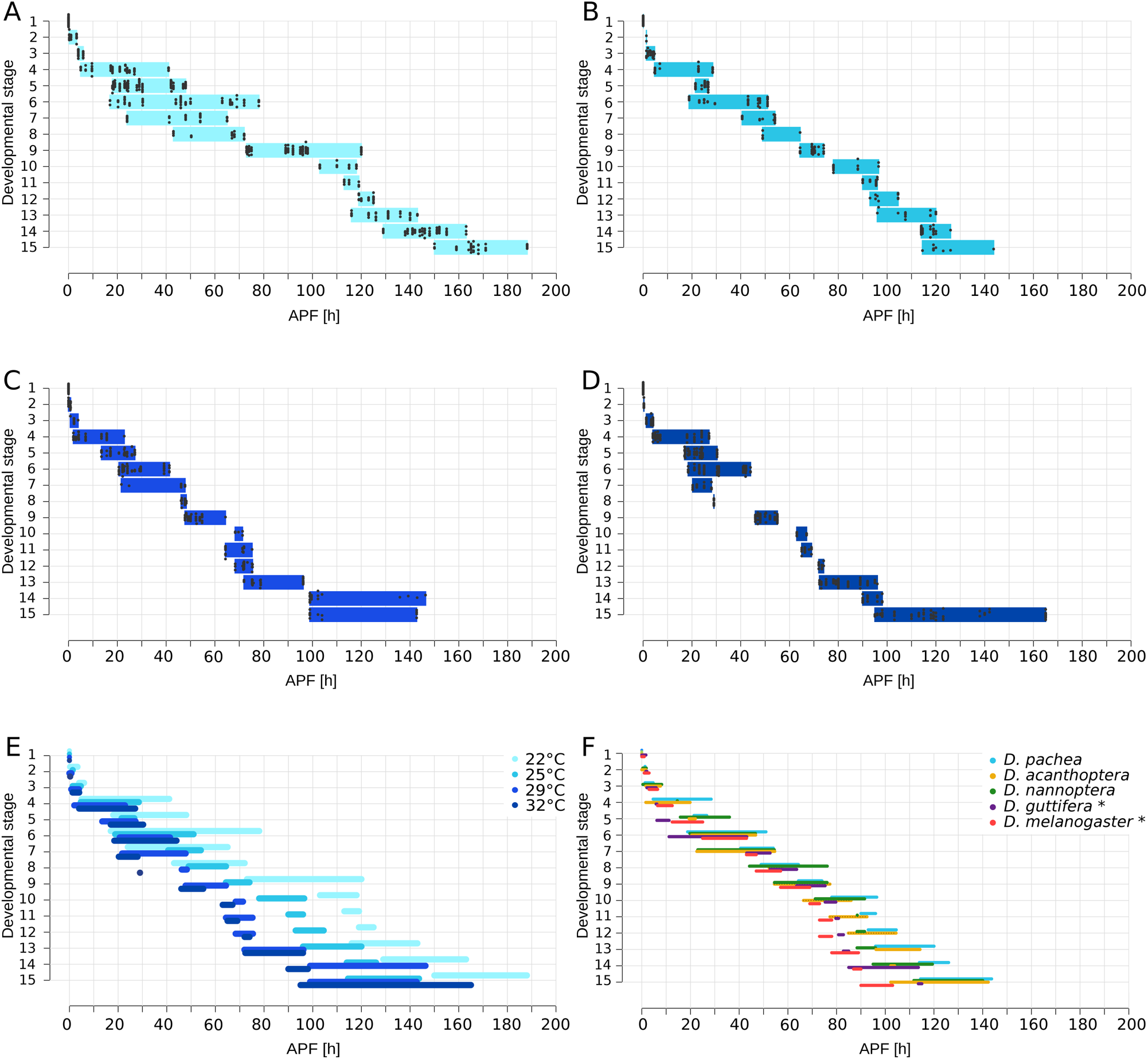
Progress of pupal development. A-D: Durations of developmental stages in *D. pachea* pupae at 22°C (A), 25°C (B), 29°C (C) and 32°C (D), observed at various time points. Temperatures are highlighted in blue colour tones according to the legend. Black dots indicate single observations (Dataset S3). E: Overlay of durations from panels A - D. F: Comparison of pupal development at 25°C in *D. pachea* (blue), *D. acanthoptera* (green) and *D. nannoptera* (yellow) based on observations of synchronized cohorts. The stages 8 to 12 were determined in *D. acanthoptera* by time-lapse imaging of developing pupae, after removal of the anterior part of the pupal case (dotted lines). Data of *D. melanogaster* (pink) and *D. guttifera* (purple) were retrieved from Bainbridge and Bownes (1981) and Fukutomi et al. (2017), respectively. These species were indicated by stars in the legend.

### 3.3. The timing of the pupal development is conserved up to the pharate adult stage between *D. pachea* and various Drosophila species at 25°C

The white pupa stage (see Material and Methods for details) in *D. pachea* was estimated to last for 102 min ± 41 min (mean ± standard deviation) (n=9) at 25°C. This duration has to be considered as the remaining variation of developmental progress between examined individual pupae in later timing analyses (see Materials and Methods). This duration was similar to previously reported durations for *D. melanogaster* white pupae of 80-120 min, at 25°C (Bainbridge and Bownes, 1981).

At 25°C, the pharate adult stage (stage 7, Table 2) was observed about 55 h after puparium formation and emergence of adults between 115 - 145 h after puparium formation (Figures 2A). This timing was similar to those of *D. acanthoptera* and *D. nannoptera* (Figure 3B). The developmental duration from puparium formation to pharate adult (stages 1 to 7, from 0 h APF to about 55 h APF) was also similar to those reported for *D. melanogaster* and *D. guttifera* (Figure 3B) (Bainbridge and Bownes, 1981; Fukutomi et al., 2017). However, at later pupal development durations of stages were prolonged in *D. pachea, D. nannoptera* and *D. acanthoptera* compared to *D. melanogaster* and *D. guttifera*.

The emergence of the adult fly from the pupal case (stage 15) is highly variable within *D. pachea, D. nannoptera* and *D. acanthoptera. D. pachea* adults emerge between 115 - 144 h APF, *D. nannoptera* adults between 112 - 140 h APF and *D. acanthoptera* adults between 102 h - 142 h APF. The variance of this stage was significantly different between the three species (Levene’s test: F = 3.4414, Df = 2, p = 0.03847), the stage 15 being longer in *D. acanthoptera* compared to *D. pachea* and *D. nannoptera* (Figure 3B).

## 4. Discussion

### 4.1. A possible temperature-buffering mechanism during pupal development

The trend of a decrease of developmental duration when rearing temperature increases was not observed in *D. pachea* at high temperatures. Overall, pupal development duration were similar at 25°C and 32°C, while it was prolonged at 22°C. On the contrary, the duration of the overall pupal development decreases with increasing rearing temperature in *D. melanogaster* (Ashburner and Thompson Jr, 1978; Powsner, 1935). In addition, temperature fluctuations during pupal development of *D. melanogaster* are known to either increase or decrease developmental speed (Ludwig and Cable, 1933; Petavy et al., 2001). In this species, the first 24 h of pupal development are more sensitive to temperature changes compared to the rest of the pupal stage (Ludwig and Cable, 1933; Petavy et al., 2001). While *D. melanogaster* is a cosmopolitan species that lives in a wide climate range (David and Capy, 1988), *D. pachea* is a desert species endemic of the Sonora (Heed and Kircher, 1965; Markow and O’Grady, 2005). The mean daily variations of temperature of this habitat are 18°C - 42°C in spring/summer and 6°C - 32°C in fall/winter (Gibbs et al., 2003). *D. pachea* is found in the wild throughout the year but undergoes a strong population decline during August, when the seasonal temperatures are highest (Breitmeyer and Markow, 1998). However, adult *D. pachea* are particularly resistant to high-temperatures and survive up to 44°C in the wild, while most other Drosophila species revealed a decreasing survival already at 38°C (Stratman and Markow, 1998). Thus, this species may have developed some heat resistance mechanisms, physiological and/or behavioral, that results in a certain tolerance to temperature variations and would buffer temperature changes on the developmental progress. This buffering effect could potentially be important for proper development since heat stress has been reported to increase developmental instability in various species (Kristensen et al., 2003; Nishizaki et al., 2015; Polak and Tomkins, 2013). However, the specific mechanism by which temperature affects developmental stability is not well understood (Abrieux et al., 2020; Breuker and Brakefield, 2003; Carvalho et al., 2017; Enriquez et al., 2018). Rearing at a lower temperature (< 25°C) revealed slower developmental progress, indicating that a potential buffering for colder temperatures does not exist in *D. pachea*.

Alternatively, the observed buffering phenotype may be temperature independent and could perhaps ensure the emergence of the adult fly at a particular moment of the day, such as dawn or dusk, when the environmental temperature might be most suitable for the freshly emerged individual. In the last pupal stage that corresponds to the adult emergence, we observed timing variation between individuals in *D. pachea* (up to 75 h between individuals). This variation could potentially depend on individual differences or on environmental factors that we could not control, such as the light/dark illumination cycle at the moment of adult emergence. Such circadian regulation of adult emergence has been observed in various Drosophila species (Ashburner et al., 2004; Mark et al., 2021; Powsner, 1935; Soto et al., 2018). However, the important variation in the last pupal stage is also found among individuals of the same cohort (Dataset S3 and S4). Future monitoring of the emergence of adults from various cohorts collected at different moments of the day will be necessary to test this hypothesis. Future investigations will be needed to further characterize the potential temperature buffering effect during *D. pachea* development and to test the influence of the circadian rhythm in this species. In addition, we must further assess temperature dependent pupal development in a wider range of species that live in distinct climate habitats.

### 4.2. Conservation of the overall developmental progress during early pupal stages

The detailed analysis of the timing of pupal stages revealed that the first stages 1 to 7 appear to be rather synchronous among *D. melanogaster* (Bainbridge and Bownes, 1981), *D. guttifera* (Fukutomi et al., 2017), and the three closely related species *D. pachea, D. acanthoptera* and *D. nannoptera*. Later on, pupal development appears to be more variable between species. This may indicate the existence of some developmental constraints, which are limitations of phenotypic variability due to inherent properties of the developmental system (Smith et al., 1985; Wagner, 2014). Such constraints probably act on outgrowth of adult organs from primordial structures, so-called imaginal discs, that develop throughout larval stages but undergo extensive tissue growth during pupal development up to the pharate adult stage. Thereafter, the timing of development seems to be less constrained and interspecific variations were observed. At least a part of the variation in the pupal developmental timing could be attributed to the developmental marker used. As the coloration markers are qualitative, it is hard to define precise limits of each stage (ie. eyes turn progressively from yellow to red). A solution might be to identify a combination of multiple markers for each stage or to establish gene expression markers that are known to account for specific developmental processes, as it has been recently done for eye development (Escobedo et al., 2021) or male genitalia development (Vincent et al., 2019).

### 4.3. *D. pachea* embryonic and larval developmental durations appear to be longer compared to other Drosophila species

The embryonic developmental duration at 25°C has been investigated in 11 drosophila species other than *D. pachea* (Chong et al., 2018; Crapse et al., 2021; David and Clavel, 1966; Kuntz and Eisen, 2014; Powsner, 1935) (Figure 2) and ranged from 16 h in *D. sechellia* to 25 h in *D. virilis* (Chong et al., 2018; Kuntz and Eisen, 2014) (Figure 2), which appear to be shorter compared to embryonic development of *D. pachea* at the same temperature. Interspecific variation in the duration of embryonic development might rely on genetic factors, as closely related species tend to have similar embryonic developmental durations compared to those of distantly related ones (Figure 2). Overall, sample size was rather low in our experiments and only present a rough approximation of the time range of Drosohila larval and embryonic development at a single temperature. A detailed examination would be necessary to adequately refine the duration of each developmental stage.

Larval development appeared to be longer in *D. pachea* compared to those in *D. melanogaster* (Bakker, 1959; Strasburger, 1935). However, the duration of this developmental stage has been shown to be highly variable compared to the other life stages. In particular it has been shown that larvae are very sensitive to food composition and to crowding that affect food quality and food access (Matzkin et al., 2011; Vijendravarma et al., 2013). Food quality and food access in turn prolong the larval developmental duration (Matzkin et al., 2011). This effect of food on developmental duration might also probably affect embryonic and pupal stages indirectly due to nutrient contribution from the adult and larval stages. A slower development observed in *D. pachea* raised in the lab might also be due to variations in the ecdysone metabolism. In insects, ecdysone is first provided to the embryo as maternal contribution and then directly produced by the individual (Lafont et al., 2012). However, in *D. pachea* the first metabolic step of the ecdysone biosynthesis is different compared to other insect species, the conversion of cholesterol into 7-dehydrocholesterol being abolished (Lang et al., 2012). Instead, *D. pachea* metabolizes sterols produced by the Senita cactus on which they feed, such as lathosterol, and potentially campestenol and schottenol (Heed and Kircher, 1965), into steroid hormones differing in their side residues (Lang et al., 2012). Therefore, in the wild, *D. pachea* may produce different variants of ecdysone that may also differently affect developmental timing compared to the lab conditions that only provide the single ecdysone precursor 7-dehydrocholesterol. Thus, it would be interesting to compare developmental durations of *D. pachea* fed with standard *D. pachea* food used in the lab or with their natural host plant, the Senita cactus. In addition, further investigations would be needed to elucidate how temperature modulates these mechanisms.

### 4.5. Conclusion

We investigated the effect of temperature on developmental speed in *D. pachea*, a desert species. We characterized the timing of the life-cycle in this species and observed prolonged developmental durations compared to other Drosophila species. The global developmental duration during metamorphosis is similar at rearing temperatures between 25°C and 32°C although stage specific timing differences were observed. These observations indicate that *D. pachea* might potentially have evolved mechanisms to buffer the effect of temperature on developmental speed. Such mechanisms might be of importance to preserve the fitness of individuals exposed to extreme temperatures and important temperature variations during their development.

## Acknowledgements

We thank all team members of the Evolution and Genetics team, including Jean David, for stimulating and constructive discussions. We thank Virginie Courtier for comments on the manuscript and for covering a part of the experimental costs.

## Supplementary data

**Figure S1: Mouth hook morphology at the three different larval instar**.

Larval mouth hook from A: first, B: second and C: third larval instar in *Drosophila pachea*. The scale bar is 10 µm.

**Figure S2: Pupal stages in *D. pachea***.

Pupal stages of *D. pachea*, based on the characterization of *D. melanogaster* by Bainbridge and Bones (1983) (Table 2). Pupae of each stage are presented in dorsal (D) and ventral (V) view. The stage (Table 2) is indicated by a number. Arrows point to relevant morphological markers: C: stage 3, dorsal trachea still visible; D: stage 4, bubbles appear and trachea not visible; E: stage 5, distal margins of wings appear; F: stage 6, yellow body visible; G: stage 7, non-pigmented eyes visible; H: stage 8, yellow eyes appear; I: stage 9, orange eyes; J: stage 10, red eyes; K: stage 11, thorax bristles visible; L: stage 12, grey wings; M: stage 13, black wings; N: stage 14, meconium visible. The scale is 100 µm.

**Movie S1: Time-lapse of embryonic development of *D. pachea* at 25 °C**.

Out of the 28 embryos, 12 completed their development up to the larva hatching. The 16 embryos that did not complete their embryonic development were excluded from the analysis.

**Movie S2: Time-lapse of pupal development of *D. acanthoptera* from 52 h APF up to the emergence of the adult at 25°C**

Out of 5 pupae, 3 completed their development up to adult emergence. The two that died during the time-lapse were excluded from analysis.

**Dataset S1: Observations of embryonic development in *D. pachea* at 25°C**

**Dataset S2: Observations of larval development in *D. pachea* at 25°C**

**Dataset S3: Pupae cohorts for developmental timing characterization**

**Dataset S4: Observations of pupal development in *D. pachea, D. acanthoptera* and *D. nannotpera***

## Availability of data and material

The movies S1 and S2 supporting the results of this article are available in the DRYAD repository, https://datadryad.org/stash/share/dfhCAtgopC4JY6qmkjK6Q_UEMmf2WSfc1gdETPPI7gk.

