## Supplementary figures and images for "Developmental timing of *Drosophila pachea* pupae is robust to temperature changes"

### Figure S1

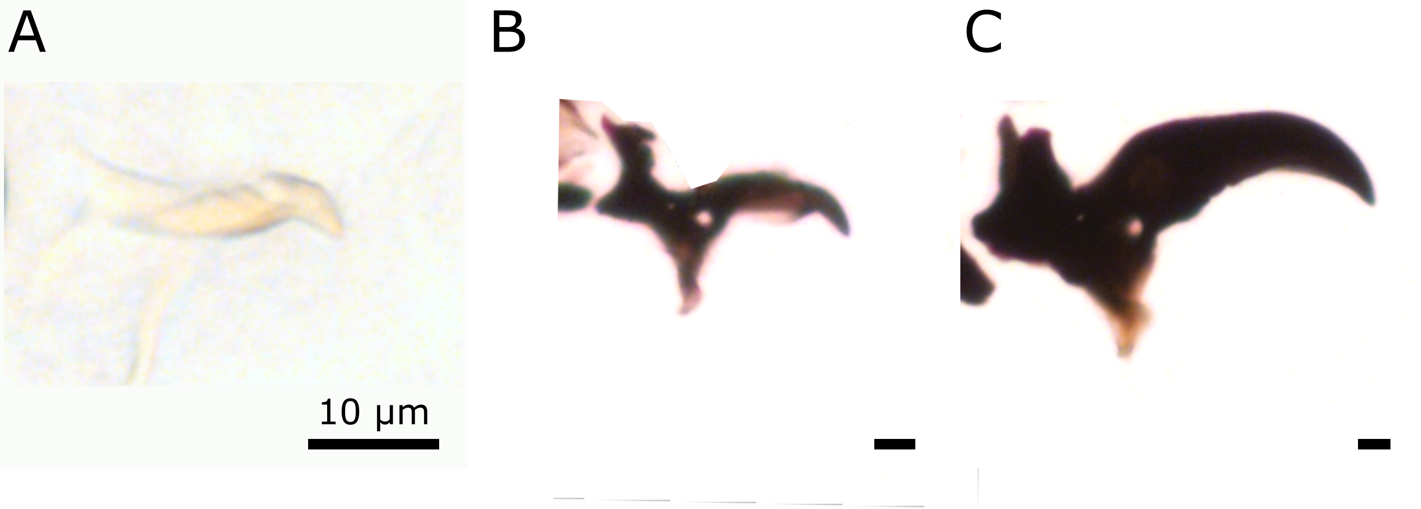

### Figure S2

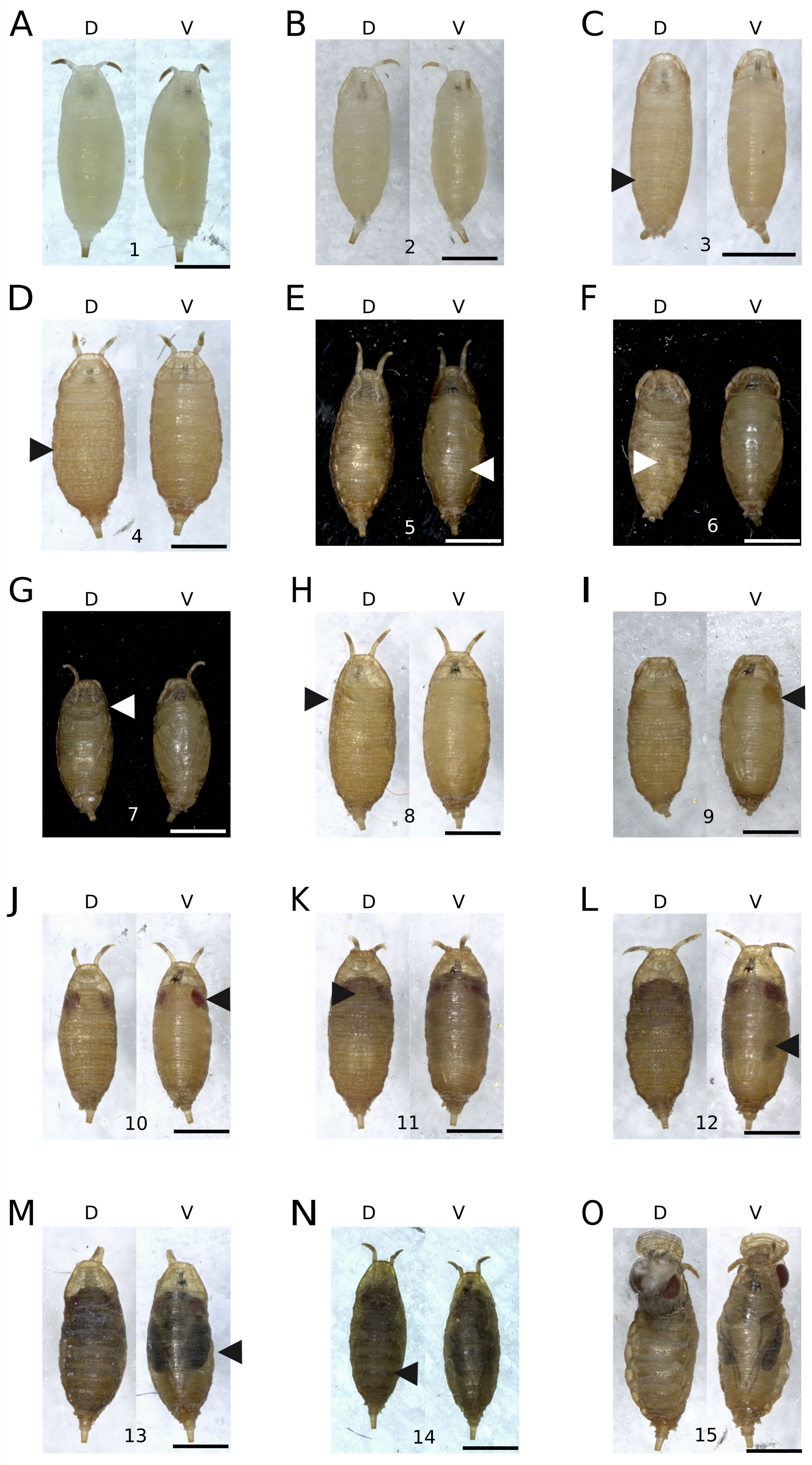
